## Supplemental Datasheet for "Exposure to live saprophytic *Leptospira* before challenge with a pathogenic serovar prevents severe leptospirosis and promotes kidney homeostasis"

**Supplementary datasheet**

Table S1: List of primary fluorochrome conjugated antibodies used in flow cytometry.

| **Marker** | **Fluorophore** | **Company** |
| --- | --- | --- |
| CD45 | Brilliant Violet 605 | Biolegend |
| CD3 | Violet Fluor 450 | TONBO biosciences |
| CD19 | Brilliant Violet 785 | Biolegend |
| CD49b | PE Dazzle 594 | Biolegend |
| CD4 | PE | Biolegend |
| CD8 | APC-Cy7 | Biolegend |
| CD44 | APC | Biolegend |
| CD62L | PE-Cy7 | Biolegend |

Figure S1

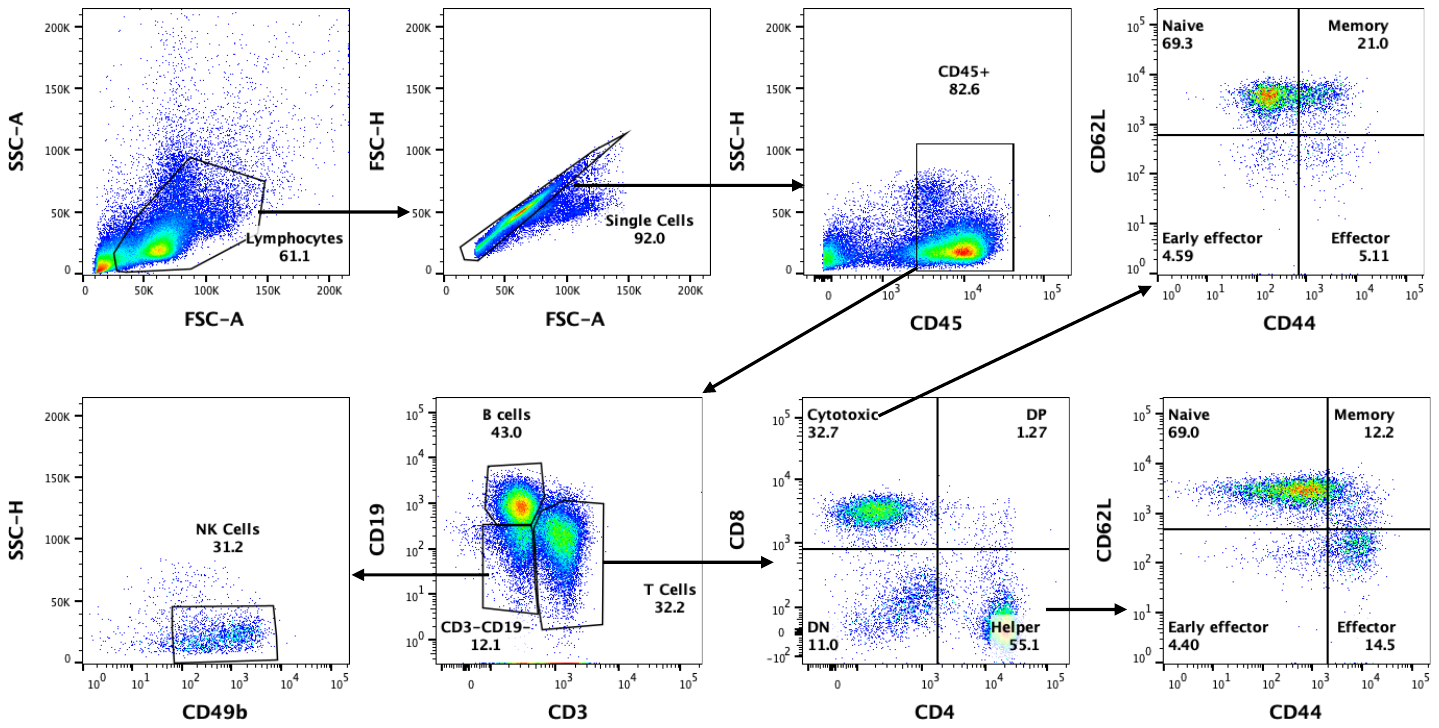

Fig S1: Gating Strategies applied for immunophenotyping of spleen. Black arrows represent the following gating events to analyze different immune cell population derived from their parent population in the spleen. X and Y axis represent specific cell surface markers to distinguish a specific cell type used in this study.

Figure S2

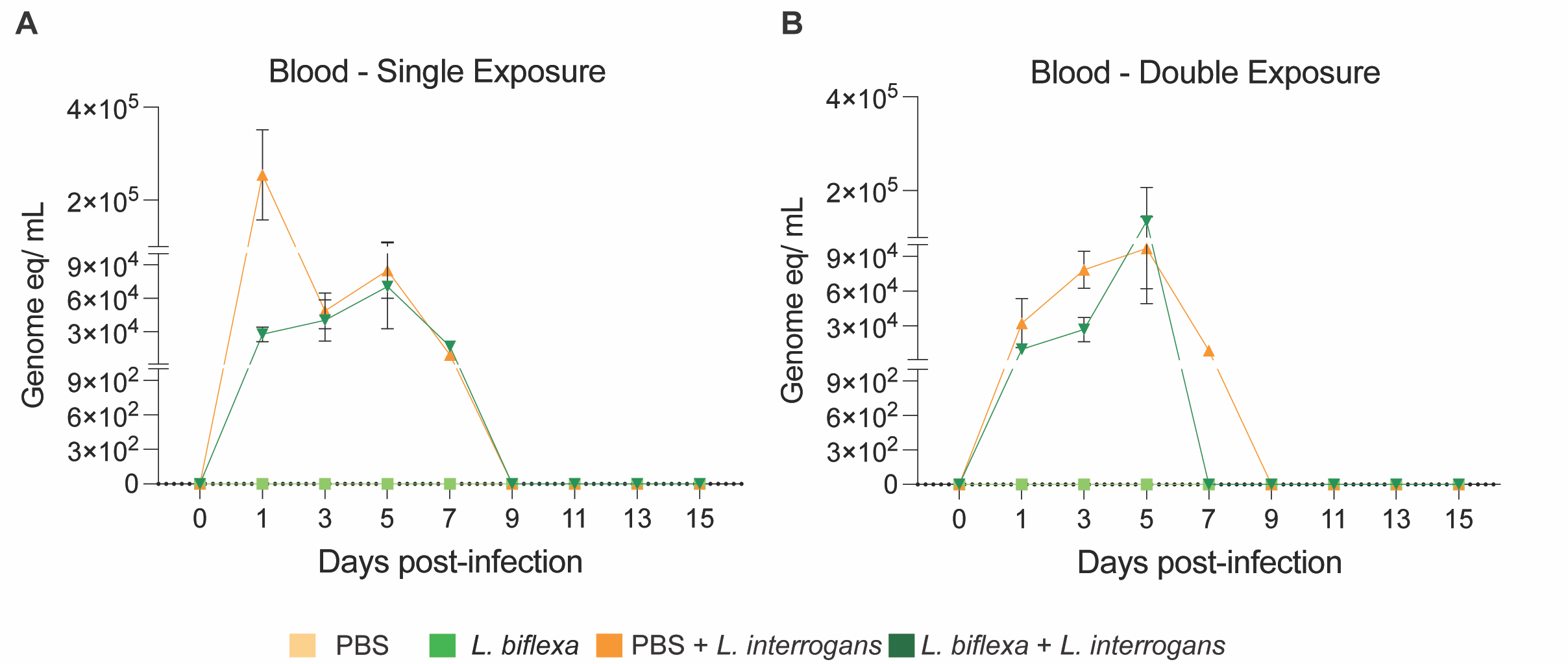

Figure S2: Real-time PCR to quantify *L. interrogans* load in blood of mice using 16s rRNA *Leptospira* specific primers and probes. Leptospiral load in A) blood from single *L. biflexa* exposure at 6 weeks followed by a subsequent *L. interrogans* challenge at 8 weeks (n=8 mice/group); and B) leptospiral load in blood from a double *L. biflexa* exposure at 6 and 8 weeks, followed by a subsequent *L. interrogans* challenge at 10 weeks (n=8 mice/group). Data represents two experiments.

Figure S3:

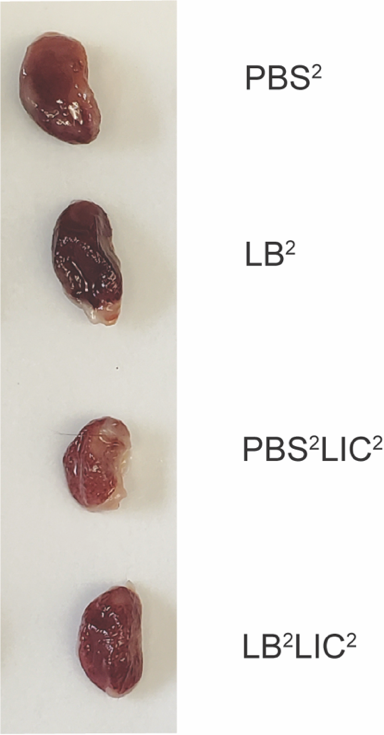

Figure S3: Morphometric analysis of kidney of mice from biweekly exposure of *L. biflexa* at 6 and 8 weeks followed by a subsequent *L. interrogans* challenge at 8 weeks. Data represents one experiment.
